# Fu Ling (*Wolfiporia cocos*) Fungal β-Glucan in Combination with *Bacillus coagulans* Improve Colitis-Related Outcomes and Cecal Fermentation in DSS- Treated Mice

**DOI:** 10.64898/2026.10.07.757334

**Authors:** Ka Lee Ma, Nelson Kei, Sui Shan Chan, Hoi Shan Kwan, Peter Chi Keung Cheung

## Abstract

Inflammatory bowel disease challenges modern gastroenterology due to high relapse rates. This study evaluated the effects of medicinal fungal sclerotium Fu Ling (*Wolfiporia cocos*), its derived β-1,3-1,6-Glucan, *Bacillus coagulans* GBI-30, 6086, and their combinations in dextran sulfate sodium-induced colitis in male BALB/c mice. The β-glucan-*B. coagulans* combination significantly improved body weight, disease activity index, colon length and architecture, mucus secretion, and cecal fermentation over single ingredient. Consistently, this combination increased total short-chain fatty acids production, particularly acetate, improved gut tight-junction organization and reduced myeloperoxidase accumulation. Shotgun metagenomic sequencing indicated that this combination reshaped the gut microbiota, notably enriching *Anaeromassilibacillus sp*., which associated with improved clinical feature. These findings highlight the potential of fungal-derived β-glucan and probiotic combinations as functional food ingredients for mitigating intestinal inflammation and supporting barrier repair.

## 1 INTRODUCTION

Inflammatory bowel disease (IBD), mainly comprising Crohn’s disease (CD) and ulcerative colitis (UC), is a chronic relapsing inflammatory disorder of the gastrointestinal tract. Although medications recommended in the 2025 American College of Gastroenterology clinical guidelines have improved clinical remission rates ^1–3^, adverse events remain common, affecting more than 60% of patients ^4^. This limitation highlights the need for safer adjunctive therapies.

A key feature of IBD pathogenesis is dysregulated mucosal immune response in genetically predisposed individuals, accompanied by pathological changes in the gut microbiota ^5^. Gut dysbiosis and altered metabolite production, including reduced short-chain fatty acid (SCFA) levels, may further exacerbate inflammation and impair immune regulation ^6, 7^. Because gut microbial metabolites support colonocyte energy metabolism and epithelial barrier integrity, interventions manipulating the gut microbiome have emerged as promising adjunctive strategies for IBD ^8–11^.

*Bacillus coagulans* is a probiotic with practical advantages for dietary intervention, as its spore-forming nature confers resistance to stomach acid, bile salts, and food processing conditions. Beyond its reported immunomodulatory effects in experimental colitis models ^12, 13^, its stability supports its incorporation into diverse food matrices. Its therapeutic potential, however, may be further enhanced when paired with a fermentable prebiotic substrate.

Fu Ling (FL), the sclerotium of *Wolfiporia cocos*, is an edible fungus with reported immunomodulatory and anti-inflammatory properties ^14–16^. Its major constituents are β-glucans, which have strong water-retention capacity due to their abundant hydroxyl groups ^17^. Our previous work showed that FL and FL-derived β-glucan are fermentable by human fecal microbiota after 24-hour *in vitro* incubation, suggesting potential prebiotic activity ^18^. However, whether combining these substrates with *B. coagulans* can enhance recovery from dextran sulfate sodium (DSS)-induced colitis remains unclear.

Therefore, the present study investigated the therapeutic potential of *Bacillus coagulans* GBI-30, 6086 (BC30), Fu Ling and FL-derived β-glucan, administered alone or in combinations, in a DSS-induced colitis mouse model. We hypothesized that the combinations would improve clinical and tissue-related indicators of colitis and would be associated with changes in gut barrier integrity and microbial fermentation.

## 2 MATERIALS AND METHODS

### 2.1 Probiotic and Prebiotic

Authentication, preparation and structural characterization of Fu Ling (FL), as well as extraction of β-glucan from FL, were performed and reported previously by our group ^18^. Briefly, fresh, unprocessed sclerotia of *Wolfiporia cocos* (Fu Ling) originating from Yunnan Province, China were obtained, washed, dried, peeled, and pulverized to a particle size of less than 0.5 mm. The FL powder was defatted with ethyl acetate and acetone, after which β-glucan was extracted overnight using sodium hydroxide. The neutralized precipitate was then purified and lyophilized to obtain β-glucan powder. FL powder used in this study was chemically characterized as over 70% of β-(1 → 3)(1 → 6)-glucans with 93% glucose monomer and low degree of branching of 0.24, whereas β-glucan powder was purified with 100% glucose monomer and a degree of branching of 0.09 ^18^. The highly stable, spore-forming probiotic *Bacillus coagulans* GBI-30, 6086 (BC30) was obtained from Ganeden Biotech (the United States). The resulting FL powder (FL) and FL-derived β-glucan powder (BG), and BC30 were stored in a desiccator at room temperature until use.

### 2.2 Animals

Male BALB/c mice (6-8 weeks, 20 g body weight) were obtained from the Laboratory Animal Services Centre, The Chinese University of Hong Kong. The mice were housed under standard laboratory conditions with a 12 h light/dark cycle at 22-25 °C and were provided free access to food and water throughout the study. The study protocol was approved by the Department of Health, Hong Kong SAR, and the Animal Experimentation Ethics Committee of The Chinese University of Hong Kong (Ref. No.: 20-218-MIS).

### 2.3 DSS-Colitis Induction and Intervention

Upon arrival, mice were fed a normal chow diet and acclimatized for 1 week. Colitis was induced by administering 4% dextran sulfate sodium (DSS) in the drinking water ^19^. Prebiotics (BG or FL; 400 mg/kg bw), probiotic (BC30; 10^9^ cfu) and their combination (BC30 combined with BG or FL) were administered daily by oral gavage ^20–22^. DSS and treatments were provided simultaneously for 6 days, after which DSS was withdrawn while treatment continued. At termination, tissue samples were collected for analyses (Figure 1).

**Figure 1.**
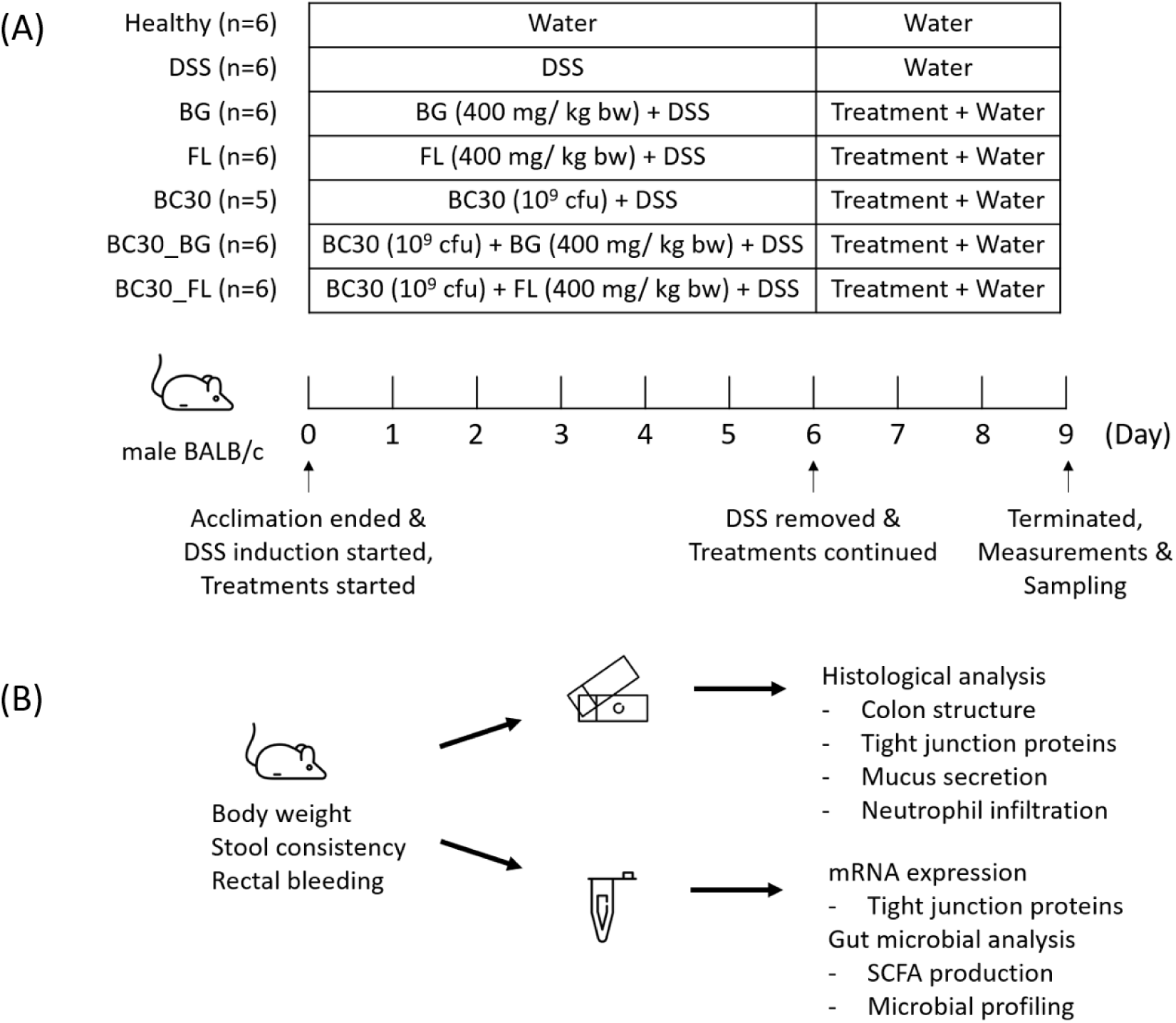
Experimental scheme for evaluating the therapeutic effects of prebiotic, probiotic and their combinations in an IBD colitis mouse model. (A) Timeline and interventions. (B) Major outcome measurements. Note: DSS, dextran sulfate sodium; BG, Fu Ling β-glucan; FL, Fu Ling; BC30, *Bacillus coagulans* GBI-30, 6086; BC30_BG, combination of BC30 and BG; BC30_FL, combination of BC30 and FL.

### 2.4 Disease Activity Index

Clinical signs of colitis were assessed using the Disease Activity Index (DAI), which comprised body weight loss, stool consistency, and gross rectal bleeding. These parameters were observed and scored daily as follows: (a) body weight loss: 0, no weight loss; 1, 1-5% weight loss; 2, 5-10% weight loss; 3, 10-15% weight loss; 4, over 15% weight loss; (b) stool consistency: 0, normal well-formed pellets; 2, pasty and semi-formed pellets which do not stick to the anus; 4, diarrhea; (c) rectal bleeding: 0, no bleeding; 2, slight visible bleeding; 4, gross bleeding. The scoring system is summarized in Table S1. The DAI was calculated as the sum of the three parameter scores, ranging from 0 (healthy) to 12 (maximal colitis activity), with slight modifications for a humane endpoint ^23^.

### 2.5 Haematoxylin and Eosin (H&E) Staining

Colon samples extending from the ascending colon to the rectum were collected and rinsed three times with phosphate-buffered saline (PBS) (pH 7.4) from the proximal to distal end. The samples were opened longitudinally and prepared as Swiss rolls, with the proximal colon positioned at the outermost layer ^24^. The rolls were fixed in 10% neutral formalin, embedded in paraffin, sectioned at 5 µm, and routinely stained with H&E. Histopathological changes were examined using Carl Zeiss PALM inverted microscope (Carl Zeiss, Germany).

### 2.6 Alcian Blue Staining

Formalin-fixed, paraffin-embedded colon sections were stained with Alcian Blue Stain Kit (pH 2.5) (G1560, Solarbio Life Sciences, China) according to the manufacturer’s protocol. Acidic mucus was stained blue, and nuclei were counterstained red.

### 2.7 Immunofluorescence Staining

Formalin-fixed, paraffin-embedded colon sections were heated in Improved Citrate Antigen Retrieval Solution (P0083, Beyotime, China) at 95 °C for 30 min to unmask antigens, followed by permeabilization with 0.025% Triton X-100 in PBS at room temperature for 10 min. The sections were then blocked with 0.1% Tween 20, 1% bovine serum albumin and 22.52 mg/mL glycine in PBS for 1 h. The slides were incubated separately with primary antibodies against ZO-1 (rabbit anti-ZO-1; ab221547, Abcam), myeloperoxidase (rabbit anti-MPO; ab208670, Abcam), claudin-1 (mouse anti-claudin-1; 37-4900, ThermoFisher Scientific, USA), followed by the corresponding Alexa Fluor 488-conjugated secondary antibodies: goat anti-rabbit IgG (ab150077, Abcam) and goat anti-mouse IgG (ab150125, Abcam). Nuclei were counterstained with DAPI using Antifade Mounting Medium with DAPI (P0131, Beyotime, China) and images were acquired using TCS SP8 Confocal Microscope System (Leica Microsystems, Germany).

### 2.8 Real-time Quantitative PCR

Segments of colon specimen were snap frozen in liquid nitrogen during termination and stored at −80°C until analysis. Specimens were ground into fine powder with mortar and pestle. Total RNA was extracted with TRIzol reagent (15596026, ThermoFisher Scientific, USA), followed by purification with Aurum^TM^ Total RNA Mini Kit (7326820, Bio-Rad, USA) to remove DSS residue ^25^, and reverse transcription with iScript III All-in-one RT SuperMix Perfect for qPCR (R333, Vazyme, China). cDNA synthesized was used as templates for RT-PCR with the aid of ChamQ SYBR Color qPCR Master Mix (Q411, Vazyme, China) on Applied Biosystems^TM^ 7500 Fast Real-Time PCR System (ThermoFisher Scientific, USA). Primers for ZO-1, occludin, claudin-1, and reference gene GAPDH were used ^26^. Primer sequences were listed in Table S2.

### 2.9 Short-Chain Fatty Acid (SCFA) Production

SCFAs were analyzed using GC-FID as previously described ^18^. Briefly, approximately 100 mg of frozen cecal content was thawed and acidified with 25% meta-phosphoric acid and 5 µL of 4-methylvaleric acid internal standard (100 mg/mL in 25% meta-phosphoric acid). The mixtures were incubated at room temperature for 30 min with continuous agitation. SCFAs were then extracted twice with 1 mL of diethyl ether and dehydrated using anhydrous sodium sulfate. A standard mixture containing acetic acid, propionic acid, butyric acid and valeric acid was prepared using the same extraction procedure. After filtration through a 0.45 µm membrane into GC vials, the SCFA samples were stored at −20 °C until analysis.

SCFAs were quantified using HP 6890 Series GC system equipped with a Quadrex 007-FFAP capillary column (30 m × 0.25 mm; i.d. 0.25 µm film) and a flame ionization detector. Five µL of each sample was injected at a split ratio of 20:1. The oven temperature was initially held at 80 °C for 3 min, increased to 100 °C at 1 °C/min and held for 5 min, then increased to 200 °C at 10 °C/min with a final hold for 4 min. Injector and detector were set to 220 °C with nitrogen as carrier gas, flowing at 1.5 ml/min.

### 2.10 Fecal Microbiome Shotgun Metagenomics Sequencing

Approximately 200 mg of frozen colon content was thawed and centrifuged to collect pellets. Total gDNA was extracted using the QIAamp Fast DNA Stool Mini Kit (51604, Qiagen, Germany), with three rounds of bead beating using sterile glass beads on Bioprep-24 Homogenizer (iGene Labserve, India). The eluted DNA was quantified using NanoDrop^TM^ One^C^ Microvolume UV-Vis Spectrophotometer (ThermoFisher Scientific, USA). Extracted DNA was dried in Savant™ SpeedVac™ DNA 120 Integrated Vacuum Concentrator System (ThermoFisher Scientific, USA) for transportation. After random fragmentation, gDNA fragments were end-repaired, indexed, amplified and sequenced on Illumina NovaSeq 6000 (PE150) according to the manufacturer’s protocols.

Image data were base-called and converted to FASTQ format using Bcl2fastq (v2.17.1.14) after preliminary quality control. Cutadapt (v1.9.1) was used to remove primers, adapters, and terminal bases with quality score below 20 to generate clean data. Host sequence contamination was removed using BWA (v0.7.12). The optimized data was processed by MEGAHIT (v1.1.3) for metagenome assembly. Assembly for each sample was performed using K-mer (79,99,119,141) and scaffold N50. The datasets generated during the current study were made available in the NCBI SRA.

Coding genes were predicted using Prodigal (v3.02), a metagenomic gene prediction tool. Redundancy among predicted gene sequences was reduced using MMseq2 to generate a unigene set with 95% identity and coverage. Gene abundance in each sample was calculated by aligning clean reads to the non-redundant unigene set by SoapAligner (v2.21), followed by functional and taxonomic annotation.

Predicted protein sequences were analyzed using BLAST (v2.2.31) against the NR (Non-Redundant Protein) ^27^, KEGG (Kyoto Encyclopedia of Genes and Genomes) ^28^, and CAZy (Carbohydrate-active enzyme) ^29^ databases, with an E-value threshold of 1e-5. DIAMOND was used to assess microbial composition against NR database, and relative abundance was generated for each sample.

Linear discriminant analysis effect size (LEfSe) analysis was performed using Galaxy ^30^, with the logarithmic LDA score threshold set at >2.0. Principal coordinate analysis (PCoA), bar plots, and association plots were generated using RStudio version 2023.09.0 with R version 4.3.1.

### 2.11 Statistical Analysis

Statistical significance was assessed using one-way ANOVA followed by Tukey’s multiple comparisons test in GraphPad Prism 9 (GraphPad Software, La Jolla, CA, USA). A p-value of <0.05 was considered statistically significant. PERMANOVA for stool microbial β-diversity and Pearson correlation test among gut microbial species and metadata were performed in R.

## 3 RESULTS

### 3.1 Body Weight, DAI, and Colon Length

At termination, the DSS group showed significant body weight loss, increased DAI, and colon shortening compared with Healthy group (Figure 2). Compared with DSS, BC30_BG significantly reduced body weight loss and DAI, whereas BC30_FL improved DAI. Colon length was restored in the BC30, BC30_BG and BC30_FL groups.

**Figure 2.**
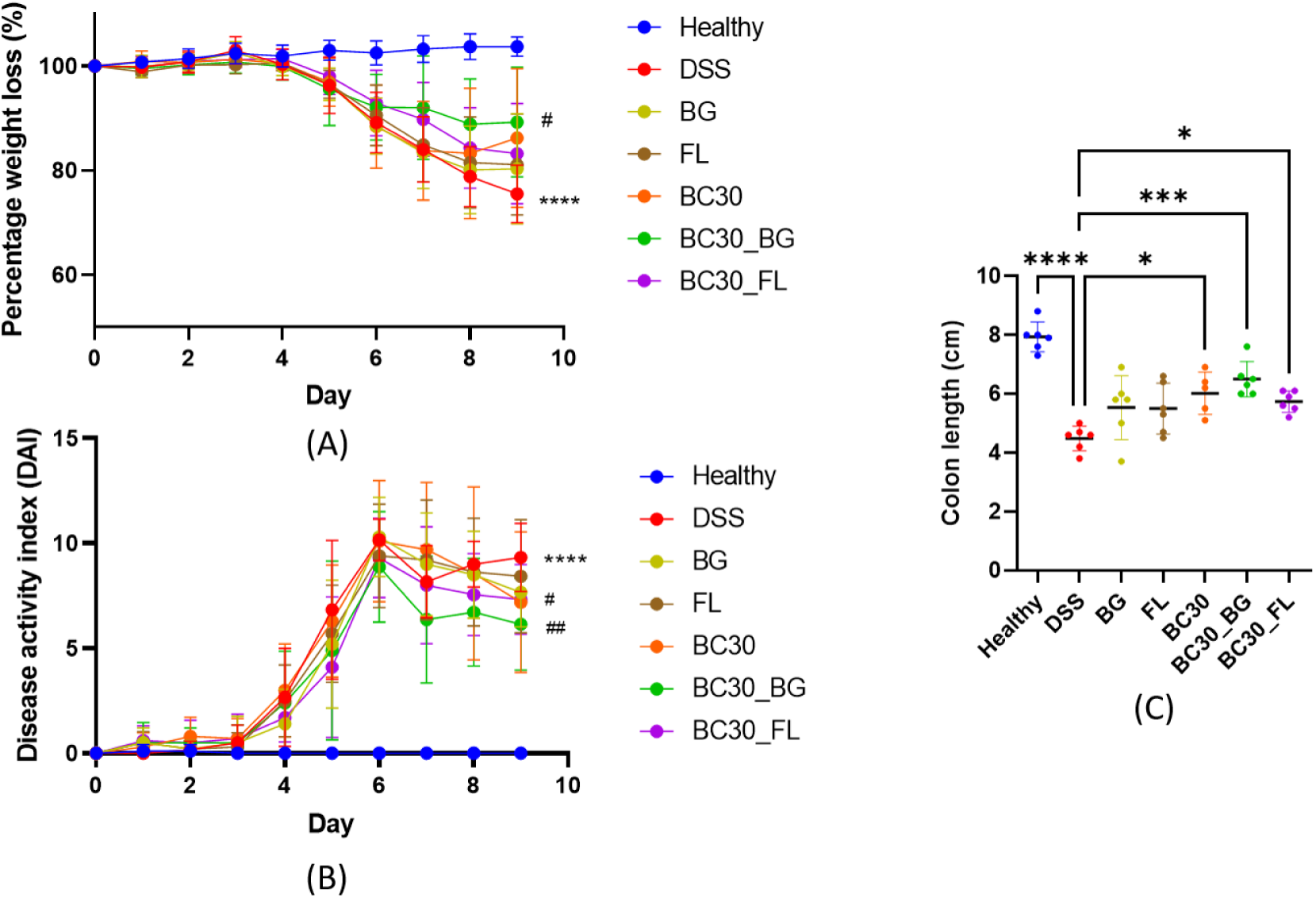
Clinical assessments of colitis severity: (A) body weight, (B) DAI and (C) colon length. Note: DSS, dextran sulfate sodium; BG, Fu Ling β-glucan; FL, Fu Ling; BC30, *Bacillus coagulans* GBI-30, 6086; BC30_BG, combination of BC30 and BG; BC30_FL, combination of BC30 and FL. N=5-6. Results are expressed as mean ± SD, analysed using one-way ANOVA, followed by post hoc Tukey’s multiple comparisons test. (A-B) **** represents p < 0.0001 compared DSS against Healthy. (A) # represents p < 0.05 compared BC30_BG against DSS. (B) # represents p < 0.05 compared BC30_FL and ## represents p < 0.01 compared BC30_BG against DSS. (C) * represents p < 0.05, *** represents p < 0.001, **** represents p < 0.0001.

### 3.2 Colon Histology and Mucus Secretion

Swiss roll sections showed that DSS-induced colitis was mainly localized to the distal colon rather than the proximal region (Figure 3A). DSS mice exhibited crypt loss, goblet cell depletion, reduced mucus secretion, and inflammatory cell infiltration into the mucosa and submucosa (Figure 3B and C). BG, BC30, BC30_BG, and BC30_FL showed better restoration of crypts and mucus-secreting goblet cells, whereas FL retained some disrupted crypts and impaired goblet cell function.

**Figure 3.**
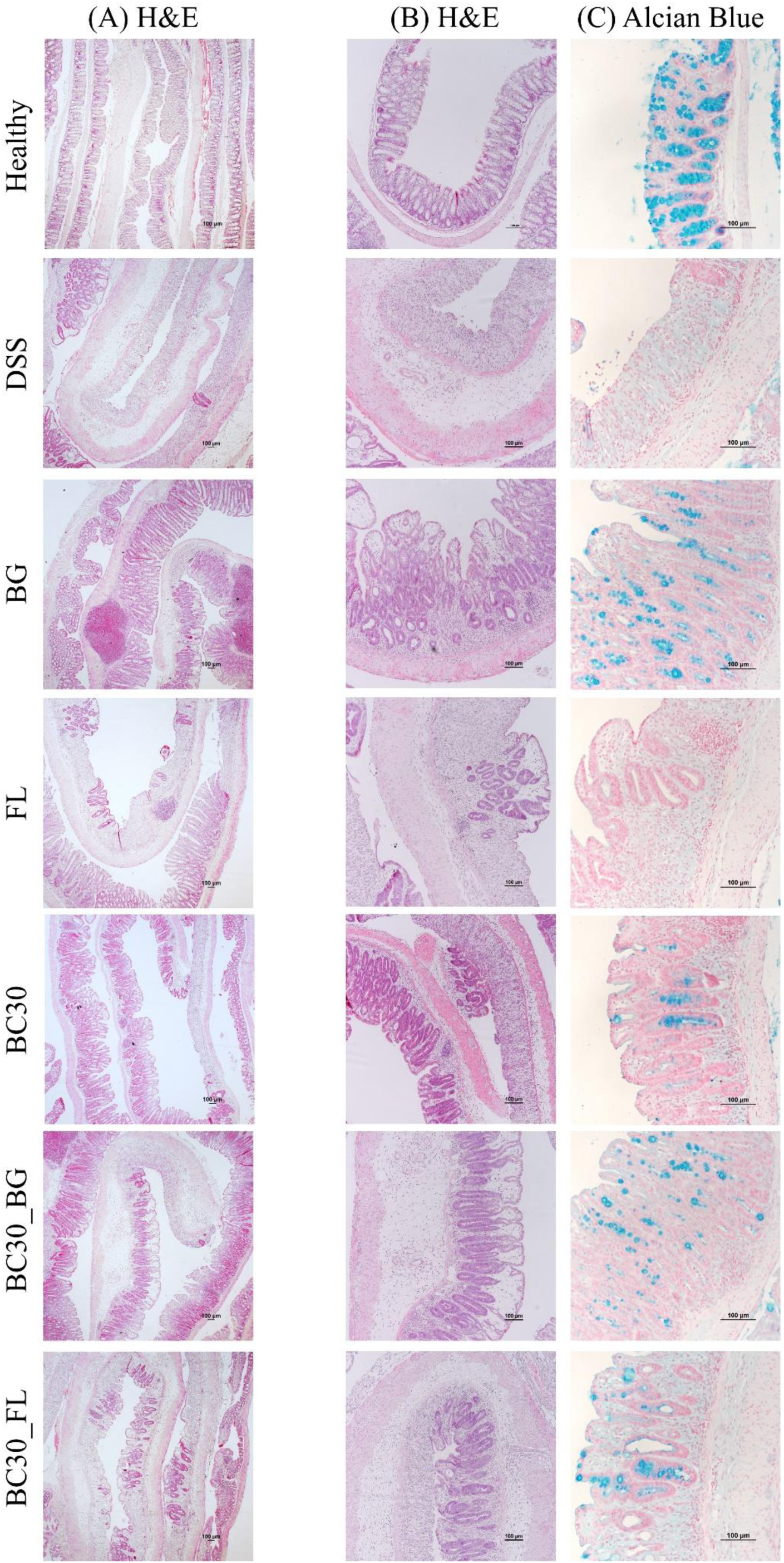
Colon histology and mucus secretion. (A) H&E staining of Swiss roll with proximal colon at the outermost layer. (B) H&E staining and (C) alcian blue/ nuclear fast red staining of distal colon. Scale bar: 100 µm. Note: DSS, dextran sulfate sodium; BG, Fu Ling β-glucan; FL, Fu Ling; BC30, *Bacillus coagulans* GBI-30, 6086; BC30_BG, combination of BC30 and BG; BC30_FL, combination of BC30 and FL.

Inflammatory cell infiltration persisted across all treatment groups but affected smaller tissue areas. Immunofluorescence staining showed increased myeloperoxidase (MPO) signals in the colon of DSS mice, which were reduced across all treatment groups (Figure 4). ZO-1 and claudin-1 organizations were disrupted in the DSS group but were restored after treatment, with a less pronounced improvement in the FL group (Figure 5). However, colon mRNA levels of tight junction proteins were not differentially expressed (Figure 6).

**Figure 4.**
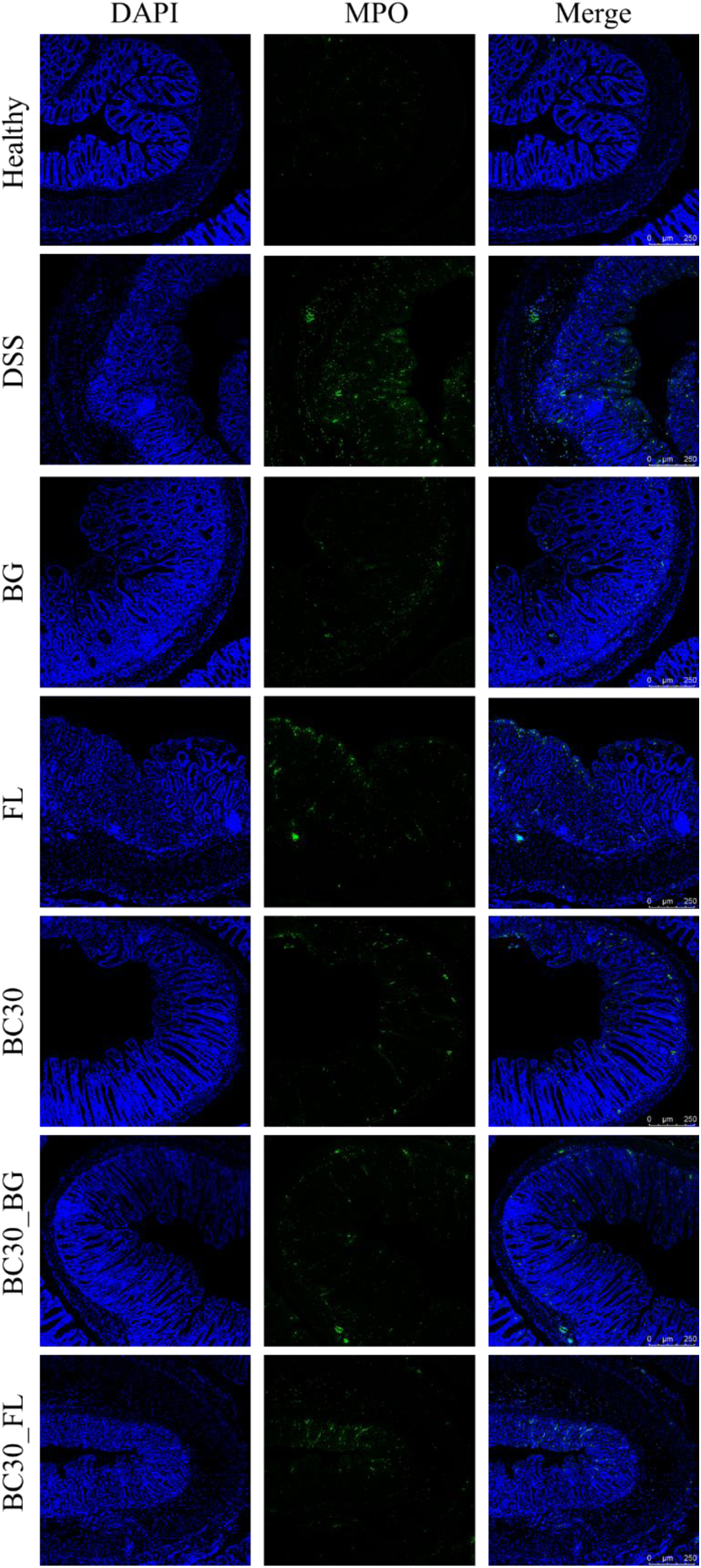
Immunofluorescence staining of colon MPO. Scale bar: 250 µm. Note: DSS, dextran sulfate sodium; BG, Fu Ling β-glucan; FL, Fu Ling; BC30, *Bacillus coagulans* GBI-30, 6086; BC30_BG, combination of BC30 and BG; BC30_FL, combination of BC30 and FL.

**Figure 5.**
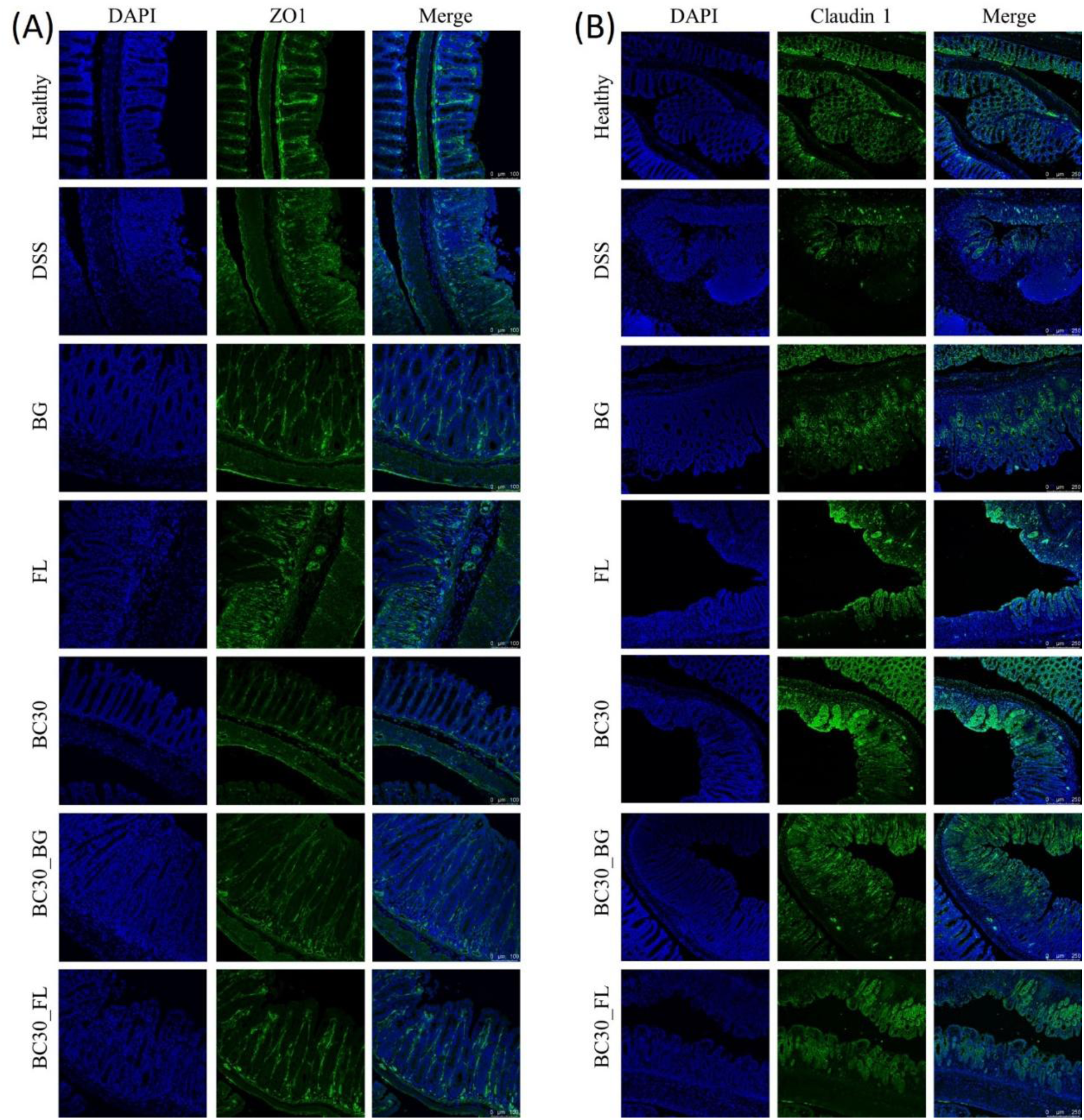
Immunofluorescence staining of colon (A) ZO-1 and (B) claudin-1. Scale bar: 250 µm. Note: DSS, dextran sulfate sodium; BG, Fu Ling β-glucan; FL, Fu Ling; BC30, *Bacillus coagulans* GBI-30, 6086; BC30_BG, combination of BC30 and BG; BC30_FL, combination of BC30 and FL.

**Figure 6.**
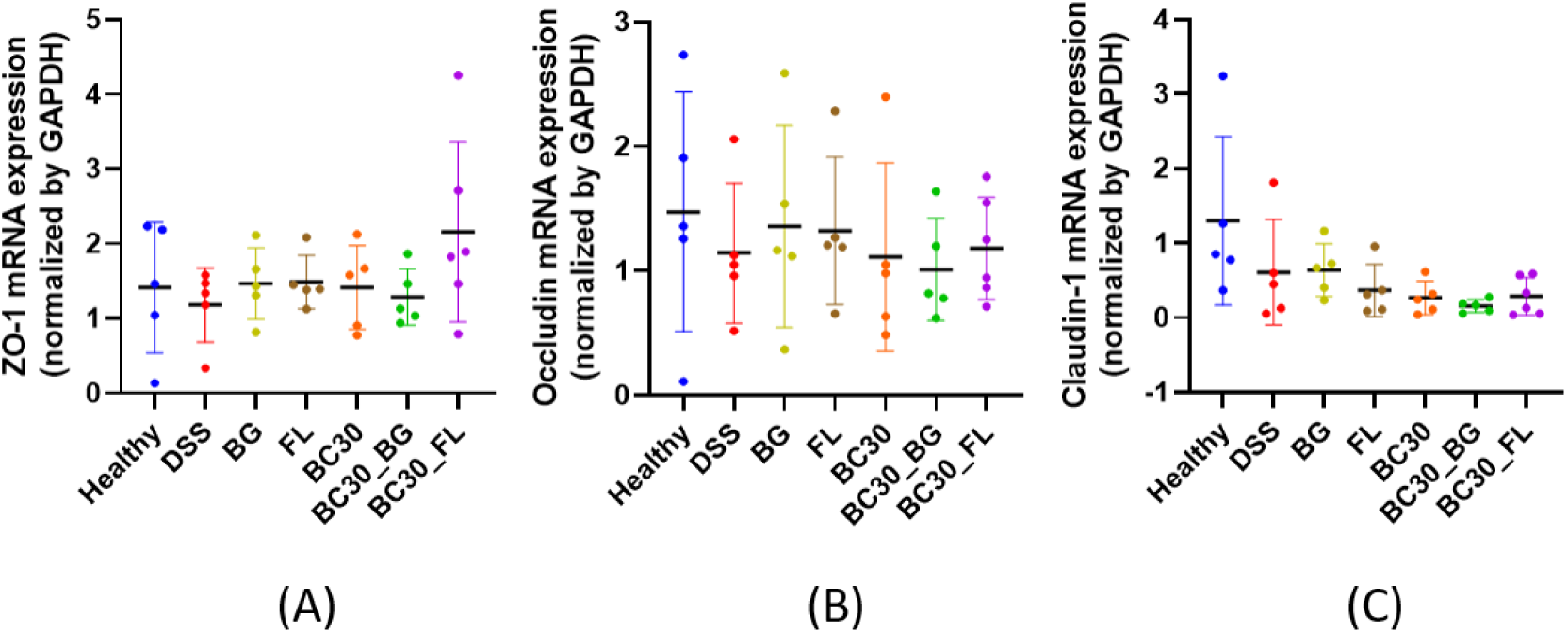
Colon mRNA levels of tight junction proteins. (A) ZO-1, (B) occludin, (C) claudin-1 mRNA level in distal colon. Note: DSS, dextran sulfate sodium; BG, Fu Ling β-glucan; FL, Fu Ling; BC30, *Bacillus coagulans* GBI-30, 6086; BC30_BG, combination of BC30 and BG; BC30_FL, combination of BC30 and FL. N=5-6. Results are expressed as mean ± SD, analysed using one-way ANOVA, followed by post hoc Tukey’s multiple comparisons test.

### 3.3 Cecal Fermentation and SCFA Production

Cecum weight and total SCFA production were significantly reduced in the DSS group but restored in the BC30 and BC30_BG groups (Figure 7). All four SCFAs were depleted in the DSS group, whereas acetic and propionic acid were replenished in the BC30 and BC30_BG groups. Valeric acid production was restored in the BC30 group.

**Figure 7.**
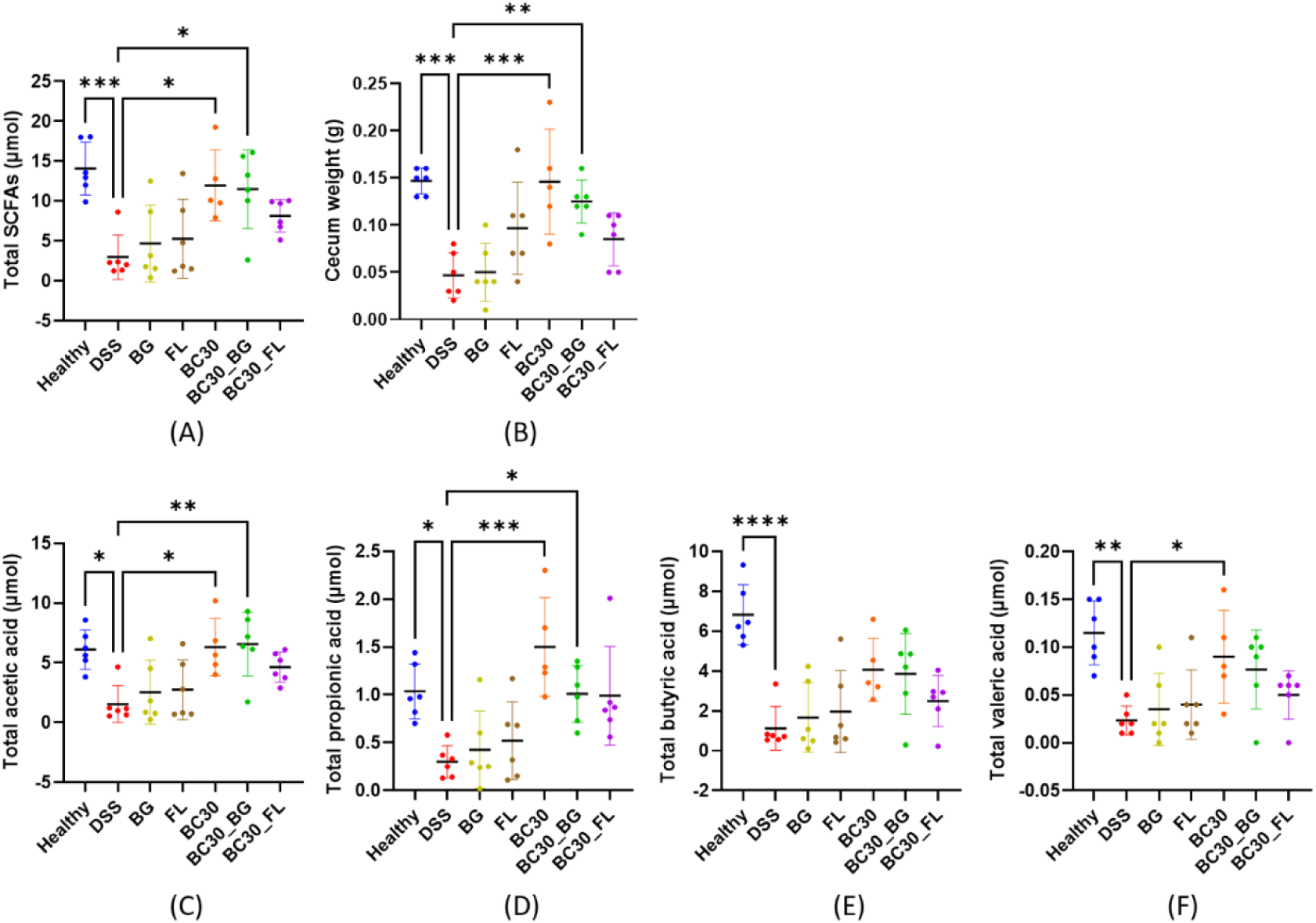
Cecal fermentation and SCFA production. (A) Total SCFA production and (B) cecum weight. Individual SCFA production including (C) acetic acid, (D) propionic acid, (E) butyric acid, (F) valeric acid. Note: DSS, dextran sulfate sodium; BG, Fu Ling β-glucan; FL, Fu Ling; BC30, *Bacillus coagulans* GBI-30, 6086; BC30_BG, combination of BC30 and BG; BC30_FL, combination of BC30 and FL. N=5-6. Results are expressed as mean ± SD, analysed using one-way ANOVA, followed by *post hoc* Tukey’s multiple comparisons test. * represents p < 0.05, ** represents p < 0.01, *** represents p < 0.001, **** represents p < 0.0001.

### 3.4 Diversity and Gut Microbial Species Enrichment

The BC30_BG group showed slightly higher α-diversity based on observed species, Chao1, ACE, Shannon, and Simpson indices. The PCoA plot showed a significant difference in β-diversity, with 38% of variance explained by group (p=0.025; Figure 8). The healthy group was clearly separated from DSS group. Although the treatments introduced variance in β-diversity, they still partially overlapped with DSS group. The relative abundance of gut microbiota at the species level is shown in Figure 9.

**Figure 8.**
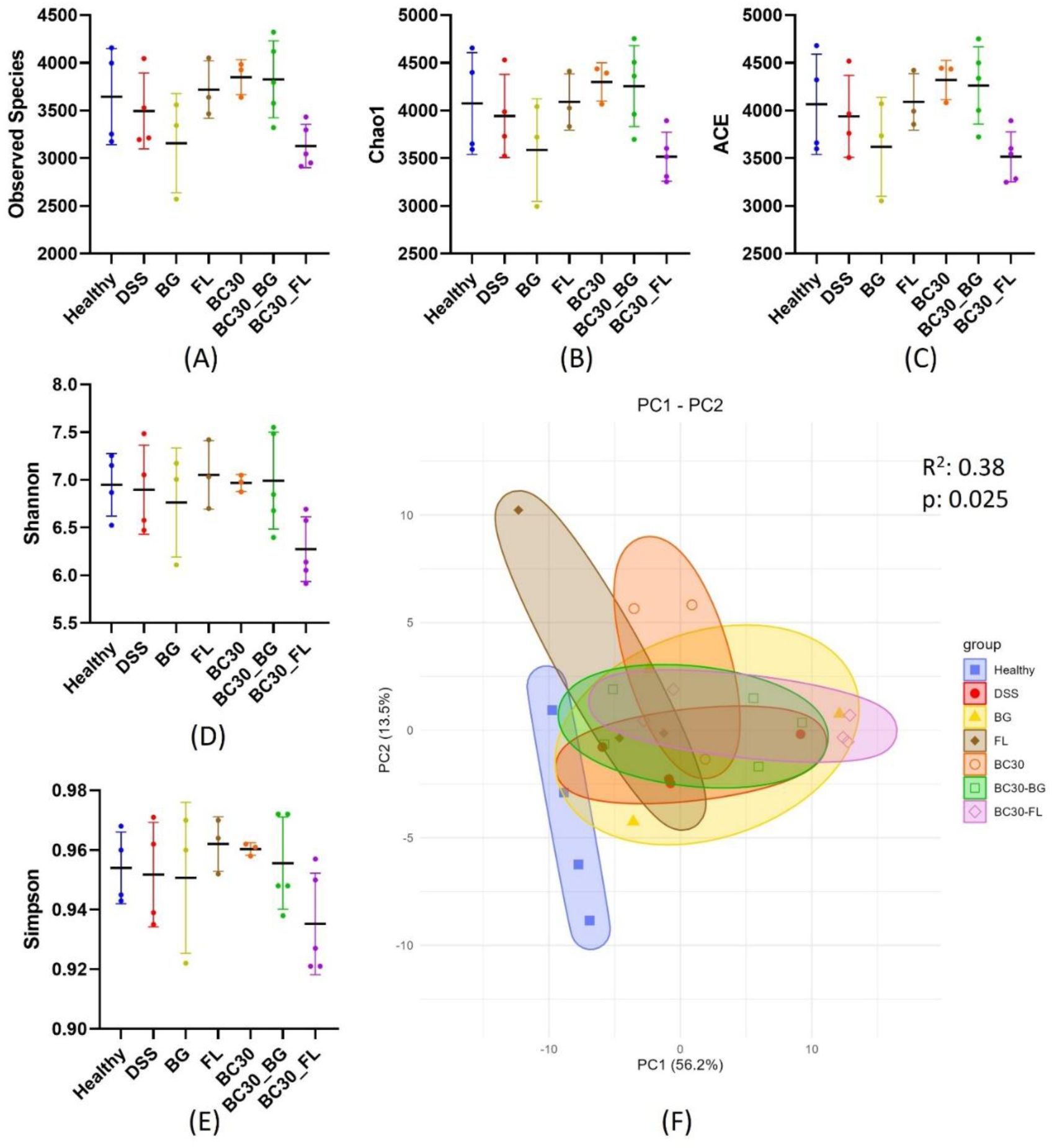
Diversity analyses of gut microbial composition. (A-E) α-Diversity indices. Results are expressed as mean ± SD, analysed using one-way ANOVA, followed by *post hoc* Tukey’s multiple comparisons test. (F) PCoA graph for β-diversity. Results are analysed using PERMANOVA. Note: DSS, dextran sulfate sodium; BG, Fu Ling β-glucan; FL, Fu Ling; BC30, *Bacillus coagulans* GBI-30, 6086; BC30_BG, combination of BC30 and BG, BC30_FL, combination of BC30 and FL. N=3-5.

**Figure 9.**
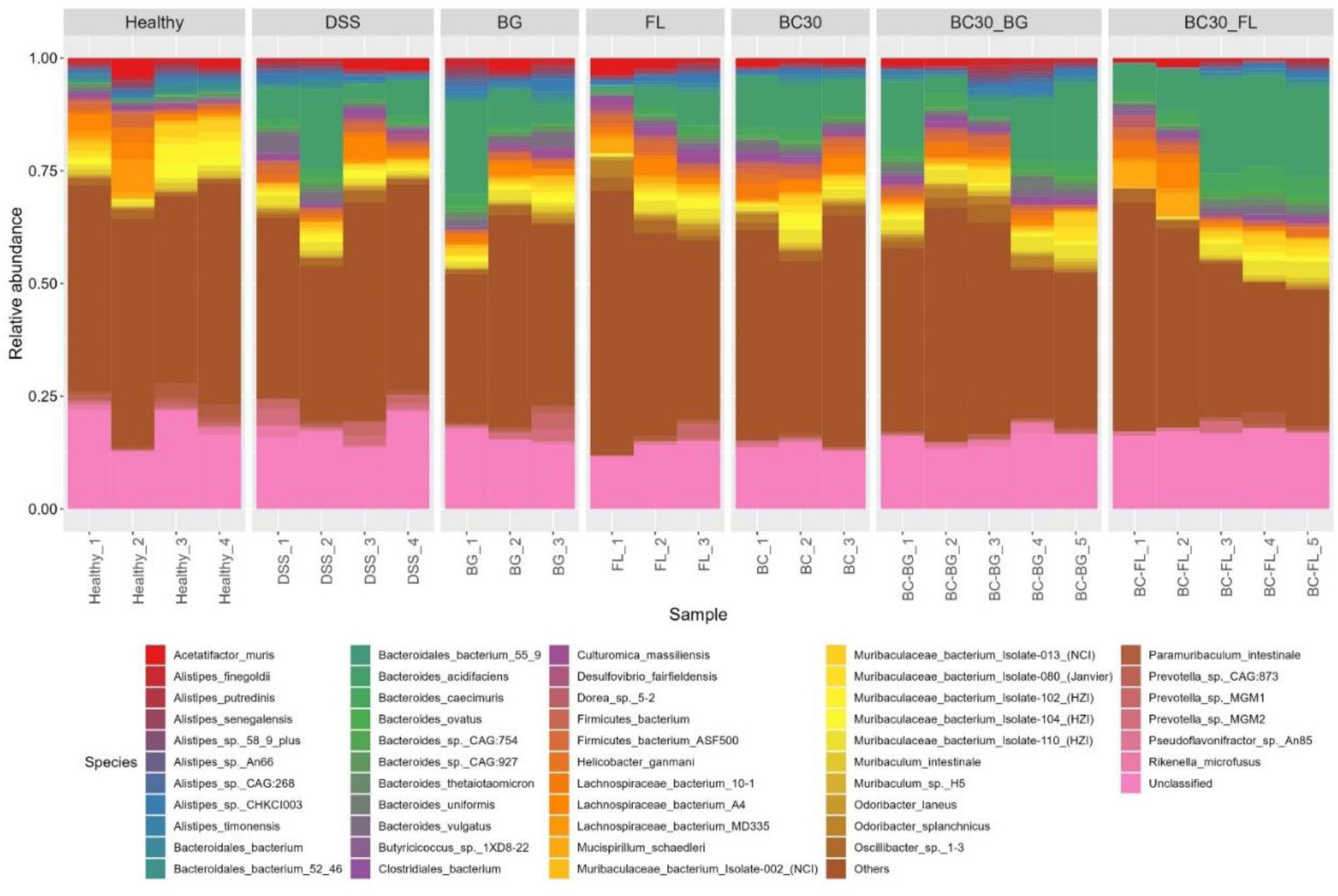
Relative abundance of gut microbiota at species level. Note: DSS, dextran sulfate sodium; BG, Fu Ling β-glucan; FL, Fu Ling; BC30, *Bacillus coagulans* GBI-30, 6086; BC30_BG, combination of BC30 and BG; BC30_FL, combination of BC30 and FL. N=3-5.

The LEfSe analysis showed a plentiful of differentially enriched species in groups. Briefly, the most predominant species were *Desulfovibrio cuneatus*, *Alloprevotella rava*, *Bacteroides vulgatus*, *Odoribacter splanchnicus*, *Bacteroides finegoldii* CAG 203, *Proteobacteria bacterium* CAG 495 and *Bacteroides acidifaciens* in the Healthy, DSS, BG, FL, BC30, BC30_BG, and BC30_FL groups, respectively (Figure 10).

**Figure 10.**
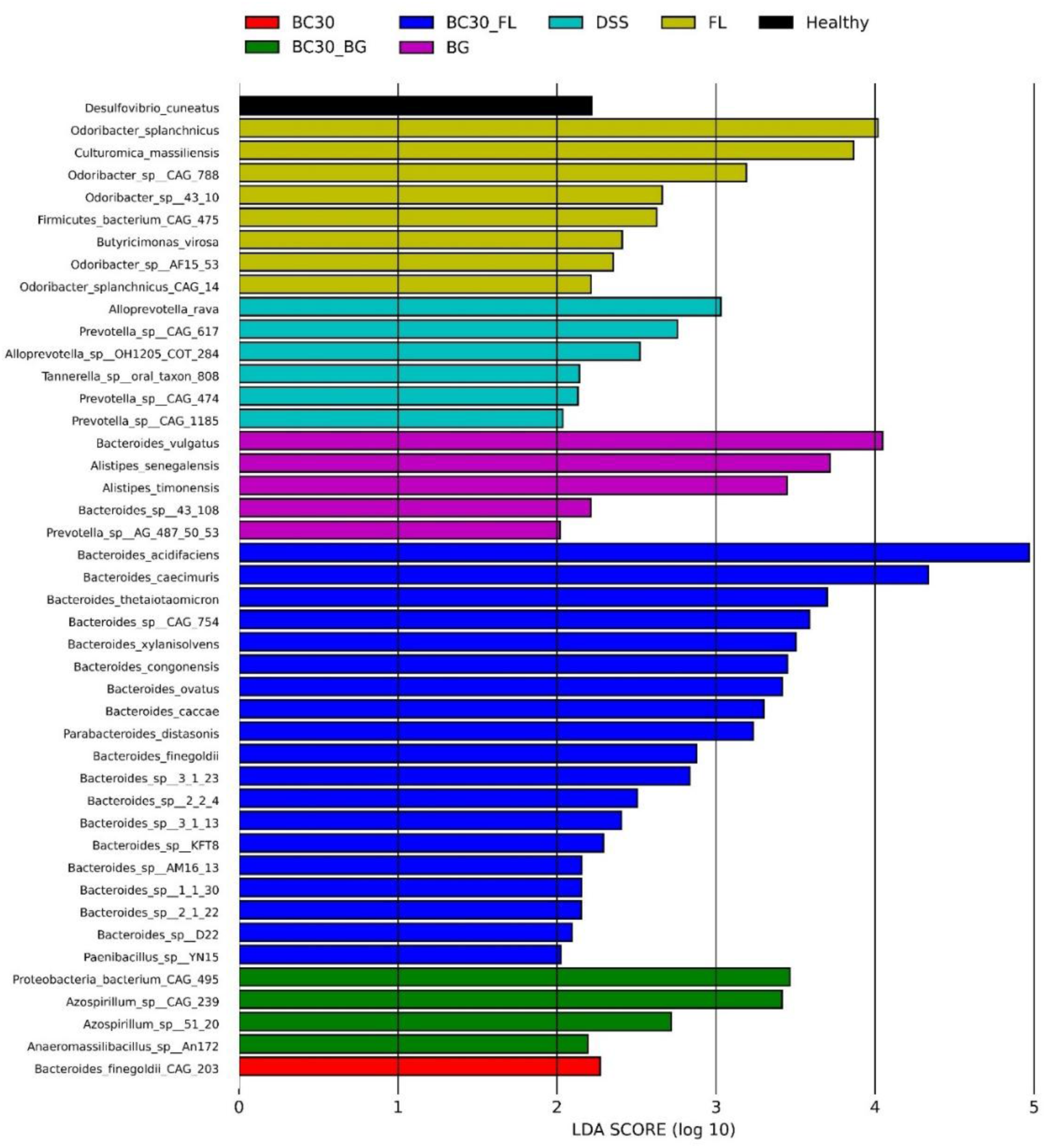
LEfSe analysis of differentially enriched microbial species found in gut microbiota. Note: DSS, dextran sulfate sodium; BG, Fu Ling β-glucan; FL, Fu Ling; BC30, *Bacillus coagulans* GBI-30, 6086; BC30_BG, combination of BC30 and BG; BC30_FL, combination of BC30 and FL. N=3-5.

### 3.5 Associations of Gut Microbial Species and Metadata

Associations between gut microbial species and mouse metadata were visualized using an association plot (Figure 11). Bacterial taxa negatively associated with DAI and claudin-1 protein or positively associated with recovery-related parameters, were considered putative recovery-associated taxa. Based on this rationale, several putative anti-colitis and pro-colitis taxa were identified. The putative anti-colitis taxa included *Anaeromassilibacillus sp.*, *Azospirillum sp.*, *Bacteroides caecimuris*, *Bacteroides sp.*, *Bacteroides xylanisolvens*, *Firmicutes bacterium*, and *Proteobacteria bacterium*. In contrast, the putative pro-colitis taxa included *Bacteroides finegoldii*, *Bacteroides vulgatus,* and *Parabacteroides distasonis*.

**Figure 11.**
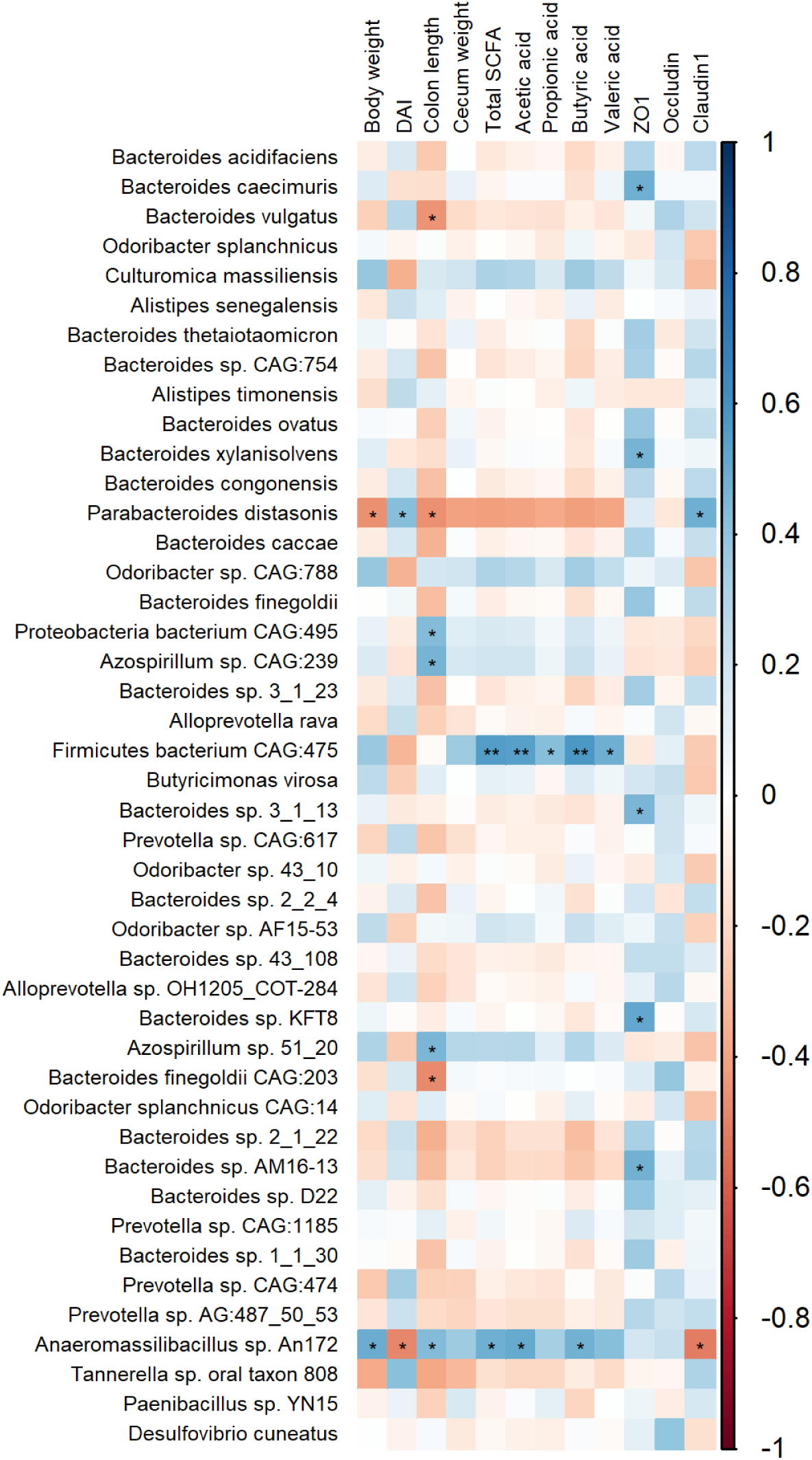
Association plot of gut microbial taxa and mouse metadata. Note: DAI, disease activity index. Results are analysed using Pearson correlation test. * represents p < 0.05, ** represents p < 0.01. Blue color represents positive correlation. Red color represents negative correlation.

### 3.6 Metabolic Pathway of Gut Microbiome

The LEfSe analysis of KEGG level 4 pathways and CAZy level 2 enzymes is shown in Figures 12 and 13. K02117 (V/A-type ATPase), K04072 (acetaldehyde/alcohol dehydrogenase, adhE), GT51 (glycosyltransferase), PL8 (polysaccharide lyase), and GT90 were differentially enriched in the BC30_BG group.

**Figure 12.**
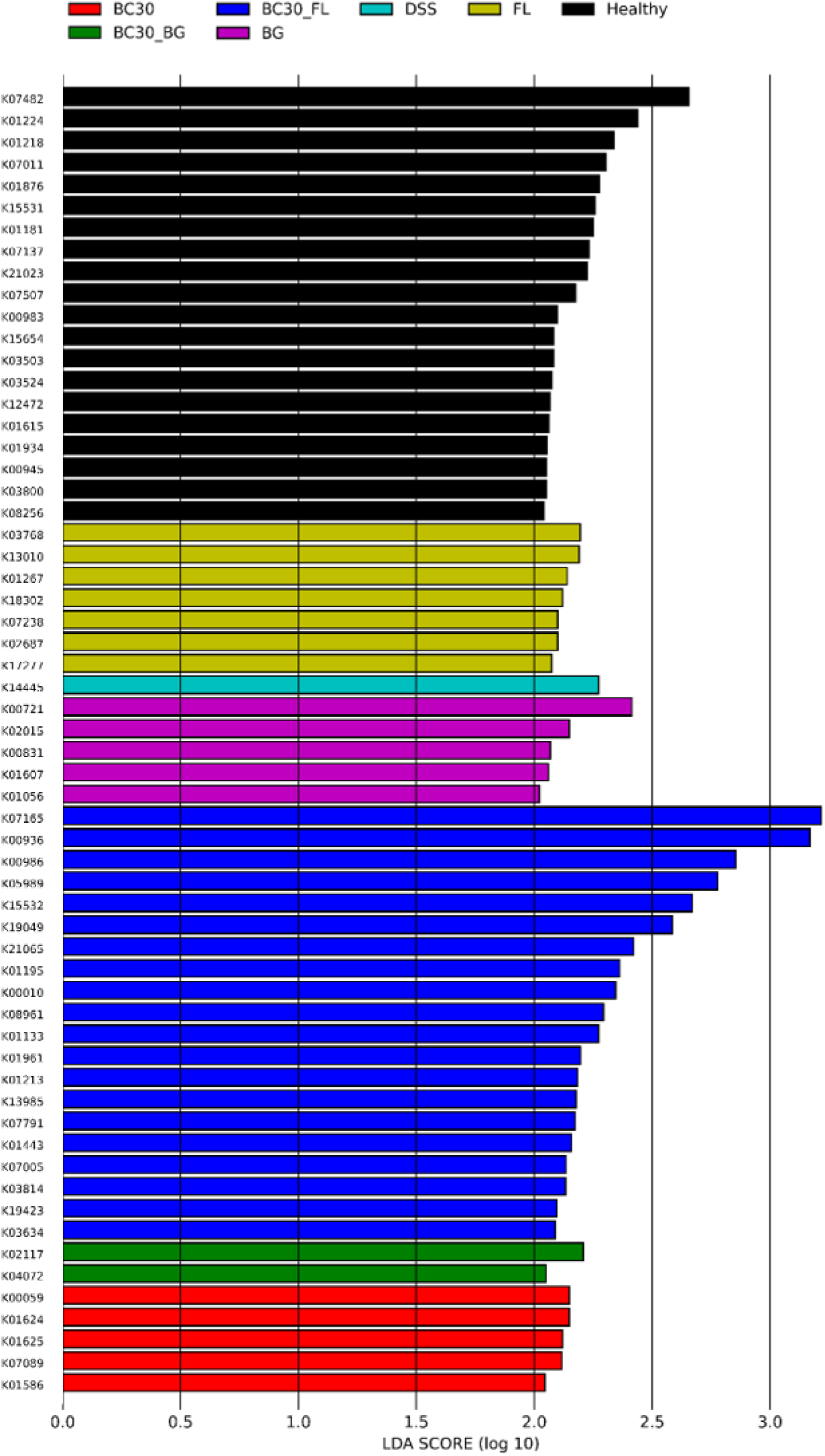
LEfSe analysis of differentially enriched gut microbial functions and pathways at KEGG level 4. Note: DSS, dextran sulfate sodium; BG, Fu Ling β-glucan; FL, Fu Ling; BC30, *Bacillus coagulans* GBI-30, 6086; BC30_BG, combination of BC30 and BG; BC30_FL, combination of BC30 and FL. N=3-5.

**Figure 13.**
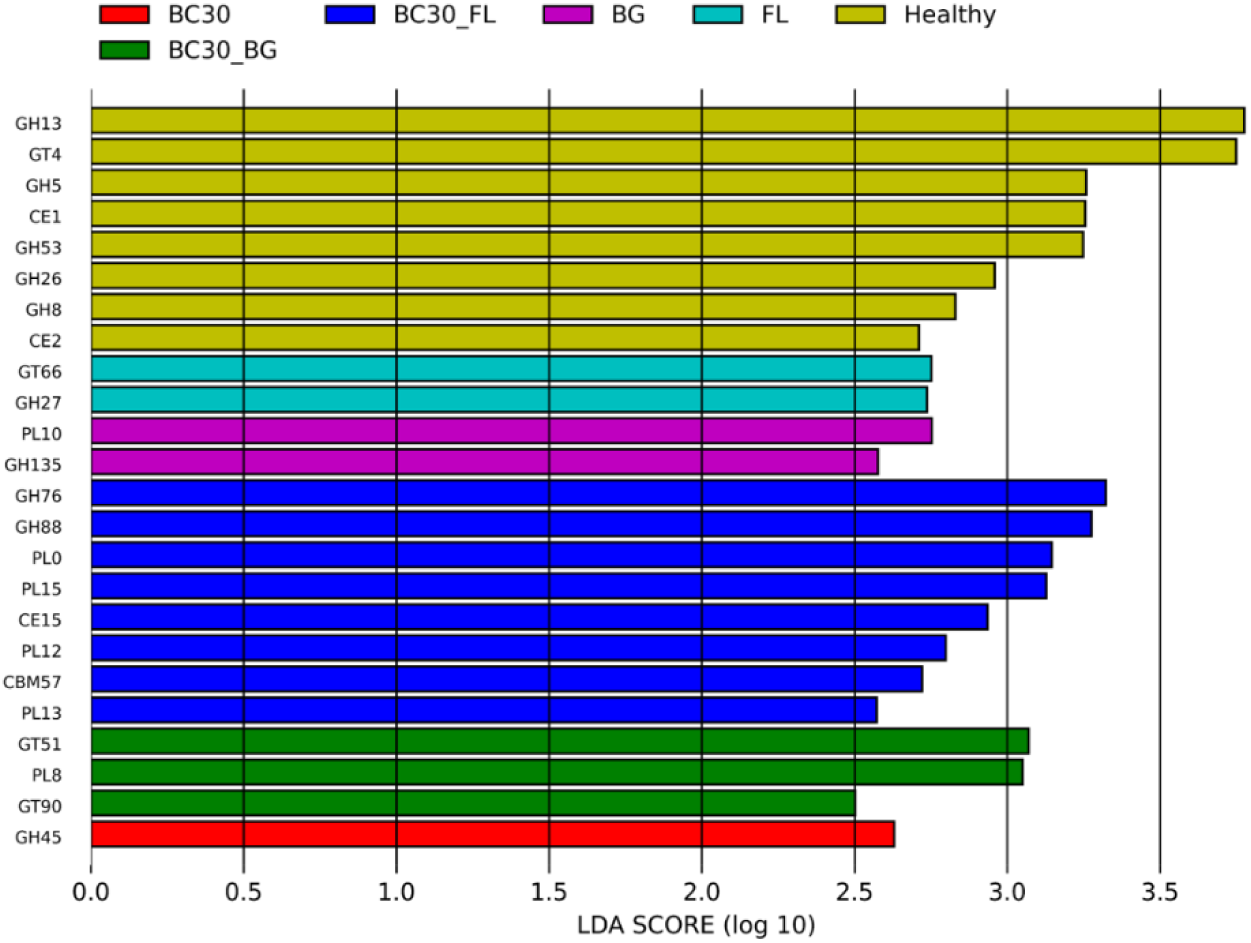
LEfSe analysis of differentially enriched gut microbial functions and enzymes at CAZy level 2. Note: BG, Fu Ling β-glucan; FL, Fu Ling; BC30, *Bacillus coagulans* GBI-30, 6086; BC30_BG, combination of BC30 and BG; BC30_FL, combination of BC30 and FL. N=3-5.

## 4 DISCUSSION

### 4.1 BC30_BG Boosted Colitis Recovery over Probiotic or Prebiotic

DSS administration in mice induces clinical and histological features that resemble human UC, making this model useful for evaluating interventions that support intestinal recovery ^31^. In the present study, Fu Ling (FL) and FL-derived β-glucan (BG), *Bacillus coagulans* GBI-30, 6086 (BC30), and their combinations (BC30_FL and BC30_BG) were compared as gut microbiota-targeted interventions. The findings did not show uniform benefits across treatments. Instead, BC30_BG produced the most consistent recovery-associated profile, with greater improvements in several outcomes than those observed with BC30 or BG alone.

The recovery-associated effects of BC30_BG were evident in the main clinical indicators of colitis severity. Although BC30_FL and BC30 produced partial improvements, BC30_BG was the only intervention that consistently improved body weight, DAI, and colon length significantly relative to the DSS group (Figure 2). A similar pattern was observed at the tissue level. While all treatments appeared to reduce some aspects of DSS-induced epithelial damage, BC30_BG showed the most pronounced improvement in colonic architecture, goblet cell recovery, and mucus secretion (Figures 3 to 5). The reduction in myeloperoxidase (MPO) staining further indicated attenuation of neutrophil infiltration and inflammation ^32^.

The comparatively strong performance of BC30_BG was also reflected in cecal fermentation, an important indicator of microbiota activity as the cecum serves as a major fermentation site and microbial reservoir in the intestine (Brown et al., 2018). DSS markedly suppressed fermentation and SCFA production, whereas BC30_BG was associated with restoration of total SCFAs, particularly acetic acid and propionic acid (Figure 7). These changes suggest that BC30_BG may support recovery not only by improving clinical and epithelial outcomes, but also by restoring microbial metabolic activity ^33, 34^. However, the mechanism underlying this response remains uncertain.

Genome-based CAZy annotation of BC30 identified putative GH1 and GH3 glycoside hydrolases, including annotations related to 6-phospho-β-glucosidase and β-glucosidase activity, as well as phosphotransferase system that may support the uptake of small β-glucoside-containing substrates. These annotations suggest that BC30 may have capacity to utilize oligosaccharides generated during the partial degradation of FL-derived β-1,3-glucan, rather than directly hydrolysing the intact polymer.

Accordingly, microbial cross-feeding may have contributed to the response observed in the BC30_BG group. However, the present data do not establish this process, because direct substrate degradation, strain-specific gene expression, enzyme activity, and metabolite transfer were not measured.

Overall, BC30_BG showed the most consistent improvements across clinical severity, epithelial repair, inflammatory cell infiltration and microbial fermentation. These findings support further investigation of BC30 combined with FL-derived β-glucan as a gut microbiota-targeted support in UC. Given the spore-forming stability of BC30, this combination may also have practical advantages for incorporation into food matrices, potentially broadening its applicability for IBD patients.

### 4.2 MPO Clearance and Tight Junction Restoration

MPO released by activated neutrophils contributes to pathogen clearance, but persistent MPO activity and delayed neutrophil resolution can promote recurrent inflammation and tissue damage in IBD ^35^. Elevated MPO levels in plasma and feces have been associated with disease severity in IBD patients ^36, 37^, while MPO inhibition has been shown to reduce oxidative injury and improve colonic damage in DSS-induced colitis mice ^38^. In this study, reduced MPO intensity may therefore indicate neutrophil clearance and the initiation of inflammation resolution (Figure 4). This inflammatory improvement was accompanied by changes in tight junction organization. Although quantities of colonic tight junction proteins were not markedly altered as shown in mRNA expression and fluorescence signals, DSS disrupted the distribution of ZO-1 and claudin-1, whereas treatment groups showed a more organized tight junction pattern (Figures 5 and 6). These findings suggest that DSS-induced barrier dysfunction may be driven more by disrupted tight junction architecture and epithelial injury than by reduced tight junction abundance.

ZO-1 is essential for tight junction assembly, and its disruption has been linked to increased leak-pathway permeability ^39, 40^. Claudin-1, together with other claudin family members and occludin, contributes to pore-pathway regulation ^41^. The discontinuous localization of ZO-1 and claudin-1 in DSS group may reflect loss of coordinated tight junction regulation and increased intestinal permeability, potentially allowing luminal antigens or bacteria to cross the damaged epithelial barrier. By contrast, the more continuous tight junction distribution observed after treatments in the study suggests barrier reorganization alongside mucosal healing. As increased permeability is shown to precede IBD onset or relapse ^42^. Interventions that support regulated permeability may be relevant for prevention. However, during active colitis, recovery is likely to depend more directly on inflammation resolution and wound healing.

### 4.3 Microbial Acetic Acid Production as a Potential Mediator of Colitis Recovery

Cecum is the main site of fermentation and a reservoir of fermentation products, SCFAs. DSS reduced cecum weight, suggesting suppressed fermentation, whereas BC30 and BC30_BG restored fermentation and increased total SCFAs, particularly acetic acid and propionic acid (Figure 7). This finding is consistent with previous DSS-colitis studies reporting acetic acid depletion ^43, 44^. Acetic acid may support intestinal recovery by reducing proinflammatory cytokine, enhancing hypoxia-inducible factor 1α and mucin expression in UC-derived epithelial cells ^9^. This finding supports microbial SCFA production is beneficial for preventing inflammation in intestinal mucosa and strengthening tight junction-associated barrier integrity.

In addition, acetic acid has been shown to promote cytoskeleton remodelling for gut barrier repair. Nakano et al. demonstrated that wound healing and reorganization of actin cytoskeleton were promoted by acetic acid in mouse colonic epithelial cell model and rectal administration in DSS-colitis mouse, via activation of JNK and Rho signalling pathways. Such effect was specific to acetic acid, but not propionate and butyrate ^45, 46^. Together, these findings suggest that restoration of cecal fermentation, particularly acetic acid production, may contribute to barrier repair and recovery in DSS-induced colitis.

### 4.4 Implications Revealed by Gut Microbiome

#### Microbial Taxa Associated with Colitis Recovery

Gut microbial dysbiosis is closely linked to intestinal inflammation ^7^, and microbiome profiling in colitis model can help identify microbial features associated with disease recovery. In this study, association plot and LEfSe analysis highlighted *Anaeromassilibacillus sp*. An172 as a recovery-associated taxon. Its relative abundance was positively associated with body weight, colon length, total SCFA, acetic acid and butyric acid production, and negatively associated with DAI and claudin-1, a leaky tight junction protein (Figure 11). This taxon was enriched in the BC30_BG group, which showed the most consistent recovery-related phenotype (Figure 10).

However, these taxonomic findings should be interpreted with caution, particularly for uncultured, poorly characterized or recently described organisms. The annotated species should be regarded as putative annotation rather than definitive strain-level identifications.

*A. sp*. was recently re-classified to a newly defined genus *Limousia* ^47^. It has been reported to encode multiple mucinolysomes involved in mucin utilization ^48^. Although mucus degradation is often viewed as potentially detrimental, mucin-utilizing bacteria such as *Akkermansia muciniphila* have shown beneficial effects in DSS-induced colitis mice ^49^, suggesting that controlled mucin turnover may support repair of the damaged mucosal barrier. *Anaeromassilibacillus sp*. has also been linked to colonic homeostasis through suppression of genes involved in antigen processing and presentation, as well as Th2-mediated cytokine responses ^50^.

Other taxa identified by association plot and LEfSe analysis may further contribute to the recovery-associated microbial profile. *Firmicutes bacterium* was associated with production of the main SCFAs and has been proposed to influence inflammatory pathways through cytokine modulation ^51^. *Proteobacteria bacterium*, which was enriched in the BC30_BG group with a high LDA score, has been reported to be upregulated during UC remission ^52^ and negatively associated with a multisystem autoimmune disorder ^53^.

Overall, these taxa may represent candidate microbial markers associated with accelerated recovery observed in the BC30_BG group, but their biological functions and contributions to colitis recovery require further validation. Nevertheless, these observations do not establish causality because LEfSe and correlations-based findings may be influenced by compositional effects, sample size and database annotation quality.

#### Microbial Metabolic Changes Suggested by KEGG and CAZy

KEGG and CAZy enrichment analyses suggested that the BC30_BG group may exhibit a recovery-associated functional shift in the gut microbiome (Figures 12 and 13). At KEGG level 4, LEfSe analysis identified enrichment of K02117 (V/A-type ATPase subunit) and K04072 (acetaldehyde/alcohol dehydrogenase, AdhE).

These pathways may reflect changes in microbial adaptation to the inflamed gut environment because intestinal inflammation can alter local pH and nutrient availability ^54, 55^. The enrichment of V/A-type ATPase and AdhE may be consistent with altered acid tolerance, proton handling, and fermentation-related metabolism.

Similarly, CAZy level 2 enrichment of GT51 and PL8 may suggest changes in microbial growth and polysaccharide utilization capacity, respectively. GT51 is associated with peptidoglycan biosynthesis ^56^, while PL8 is involved in the degradation of complex carbohydrates and glycosaminoglycans ^57^.

Overall, the KEGG and CAZy patterns observed in the BC30_BG group may point to a shift toward a more metabolically active microbial community. However, these annotations should be interpreted cautiously as gene enrichment does not necessarily indicate increased enzyme activity. These possibilities should be confirmed in future studies using targeted metabolomic, transcriptomic, or functional assays.

## 5 CONCLUSION

BC30_BG intervention led to pronounced recovery from DSS-induced colitis.

Its benefits were reflected in improved clinical indices and restoration of damaged colon, potentially through MPO clearance and gut microbial production of acetic acid. These may involve in inflammation resolution and tight junction protein reorganization. These findings support the utilization of gut microbiota-targeted treatments for human UC. As a heat-stable probiotic, this combination can be incorporated into diverse food matrices suitable for IBD patients, thereby extending its potential benefits to a broader patient population.

## AUTHOR CONTRIBUTIONS

Conceptualization: K.L.M.; Methodology: K.L.M.; Investigation: K.L.M.; Formal analysis: K.L.M., N.K, S.S.C.; Writing – original draft: K.L.M; Writing – review & editing: N.K., S.S.C., H.S.K., P.C.K.C.; Supervision: P.C.K.C.; Funding acquisition: P.C.K.C. All authors have read and agreed to the published version of the manuscript.

## DATA AVAILABILITY

Raw sequencing reads generated in this study are available in the NCBI Sequence Read Archive under BioProject accession PRJNA1509518.

## ACKNOWLEDGMENTS

The authors thank the technical staff of the School of Life Sciences, The Chinese University of Hong Kong, for assistance with animal experiments and laboratory analyses.

