## Supplementary Tables for "Fu Ling (*Wolfiporia cocos*) Fungal β-Glucan in Combination with *Bacillus coagulans* Improve Colitis-Related Outcomes and Cecal Fermentation in DSS- Treated Mice"

**Table S1.** Disease Activity Index (DAI) scoring parameters for colitis.

| **Body weight loss** | | **Stool consistency** | | **Rectal bleeding** | |
| --- | --- | --- | --- | --- | --- |
| 0 | no weight loss | 0 | normal well-formed pellets | 0 | no bleeding |
| 1 | 1-5% |  |  |  |  |
| 2 | 5-10% | 2 | pasty and semi-formed pellets which do not stick to the anus | 2 | slight visible bleeding |
| 3 | 10-15% |  |  |  |  |
| 4 | >15% | 4 | diarrhea | 4 | gross bleeding |

**Table S2.** Primers for RT-PCR amplification.

| **Gene** | **Primer Sequence** |
| --- | --- |
| GAPDH | F: 5’-GCAAAGTGGAGATTGTTGCCAT-3’  R: 5’- CCTTGACTGTGCCGTTGAATTT-3’ |
| ZO-1 | F: 5’-ACCCGAAACTGATGCTGTGGATAG-3’  R: 5’- AAATGGCCGGGCAGAACTTGTGTA-3’ |
| OCLN | F: 5’-AGACTACACGACAGGTGGGG-3’  R: 5’- CTGCAGACCTGCATCAAAAT-3’ |
| CLDN1 | F: 5’-TGCCCCAGTGGAAGATTTACT-3’  R: 5’- CTTTGCGAAACGCAGGACAT-3’ |

GAPDH: glyceraldehyde-3-phosphate dehydrogenase; ZO-1: zonula occludens-1; OCLN: occludin; CLDN1: claudin-1.
